# Mathematical Modelling of Reoviruses in Cancer Cell Cultures

**DOI:** 10.1101/2024.07.12.603333

**Authors:** Arwa Abdulla Baabdulla, Francisca Cristi, Maya Shmulevitz, Thomas Hillen

## Abstract

Oncolytic virotherapy has emerged as a potential cancer therapy, utilizing viruses to selectively target and replicate within cancer cells while preserving normal cells. In this paper, we investigate the oncolytic potential of unmodified reovirus T3wt relative to a mutated variant SV5. In animal cancer cell monolayer experiments it was found that SV5 was more oncolytic relative to T3wt. SV5 forms larger sized plaques on cancer cell monolayers and spreads to farther distances from the initial site of infection as compared to T3wt. Paradoxically, SV5 attaches to cancer cells less efficiently than T3wt, which lead us to hypothesize that there might be an optimal binding affinity with maximal oncolytic activity. To understand the relationship between the binding process and virus spread for T3wt and SV5, we employ mathematical modelling. A reaction-diffusion model is applied, which is fit to the available data and then validated on data that were not used for the fit. Analysis of our model shows that there is an optimal binding rate that leads to maximum viral infection of the cancer monolayer, and we estimate this value for T3wt and SV5. Moreover, we find that the viral burst size is an important parameter for viral spread, and that a combination of efficient binding and large burst sizes is a promising direction to further develop anti-cancer viruses.

## 1. Introduction

Oncolytic virotherapy has emerged as a compelling strategy for addressing the challenges of cancer treatment. By leveraging the ability of oncolytic viruses to selectively target and replicate within cancer cells, this approach aims to achieve tumor cell lysis without causing harm to healthy cells [53; 22; 31]. The consequential activation of the host immune system, promoting an anti-cancer immune response, further adds to the potential therapeutic benefits [14].

Various oncolytic viruses, including adenoviruses [25; 26; 28; 35; 40], herpes simplex virus [34; 3; 54; 55], vaccinia viruses [62; 58; 59], measles virus [7; 47], vesicular stomatitis virus (VSV) [32; 20] and reovirus [48; 60; 29; 43; 11; 12] have been investigated for their suitability in oncolytic virotherapy. Viruses are genetically modified to enhance their oncolytic potential and ensure selective targeting of cancer cells. Preclinical studies utilizing in vitro and in vivo models have been conducted to evaluate the safety and efficacy of these modified viruses [27; 37].

This research focuses on the oncolytic potential of reovirus [44; 45; 42; 64; 46]; a double-stranded RNA nonpathogenic virus with natural tropism to the enteric tract of mammals. The unmodified laboratory strain of reovirus serotype 3/T3DPL (T3wt) has demonstrated natural capabilities for infecting and lysing tumors under both in vitro and in vivo conditions [56; 48; 50; 11; 12]. T3wt is currently undergoing evaluation in over 30 clinical trials targeting various cancer types, including metastatic breast cancer [6; 50], prostate cancer [18; 57], and colorectal cancer [48; 1]. Additionally, it has progressed to phase III clinical trials as a potential therapeutic intervention for breast cancer [21; 6].

While T3wt is consistently well-tolerated [23; 24; 25; 26; 27], the majority of patients do not respond to reovirus therapy, and overall responses remain underwhelming [22]. A recent summary of oncolytic viral clinical trials as a whole similarly highlight that only ∼21% of patients show some response to oncolytic viruses [28]. Mutants of T3wt have therefore been selected to enhance oncolytic activity on cancer cells in vitro and improve tumor regression and survival in animal tumor models in vivo [11; 21; 44; 45; 42; 64]. One such mutant, SV5, demonstrated significant improvement of oncolysis in vivo that correlated with key parameters in cell culture including binding percentage, the distance of virus spread from primary sites of infection, and plaque sizes in cell monolayer experiments [11; 12]. The outcomes of these assessments underscore the pivotal importance of enhancing the distance that oncolytic viruses travel before reinfection as a critical mechanism for optimizing therapeutic interventions in the context of oncolytic virotherapy. Intriguingly, the experimental results reveal a discernible connection between the binding rate and the distance a viral infection spreads over the cell monolayer, suggesting that a reduction in the binding rate can lead to more extensive and distal viral spread, characterized by larger plaque sizes. This observation highlights a distinct advantage that the mutated SV5 virus possesses over the wild-type T3wt, despite similarities in viral production and cell death. These findings motivate the mathematical question addressed in this study, which is, what are the optimal values of virus binding rates that retain sufficient cell attachment to permit efficient infection of cells but also allow further distance of virus spread before re-infection to produce larger areas of virus dissemination?

The phenomenon of viruses exhibiting reduced binding to host cells has been extensively documented in the scientific literature. Notably, reovirus variants with diminished affinity for sialic acid have been identified in both murine and human species. A sialic-acid-binding-deficient reovirus variant exhibited heightened infectivity when compared to the wild-type reovirus in polarized epithelial cells from apical or basolateral orientations [19]. In the context of rotavirus mutants incapable of binding to sialic acid, although these mutants displayed slower replication and lower titres in mouse cancer cell lines MA104, they paradoxically exhibited increased pathogenicity in mice [63]. This underscores the nuanced relationship between viral binding capabilities and infection outcomes under specific conditions.

Expanding beyond reoviruses and rotaviruses, other viruses have demonstrated the capacity to spread more extensively in monolayer cell cultures without a concomitant increase in replication. For instance, vaccinia virus-infected cells repel superinfecting virions, resulting in enhanced viral spread [17]. Additionally, reduced adsorption rates to host bacteria have been linked to increased plaque size in phages [24]. Moreover, various virus variants of polyomavirus, parvovirus, and Sindbis virus, characterized by deficiencies in binding, have been shown to generate larger plaques in vitro [5; 52; 38; 10]. Importantly, these variants exhibited higher pathogenicity and increased spread in vivo.

Building on experimental data from [12] we address key questions through mathematical modeling: How does the viral spread distance correlate with the binding rate? What is the dependence of the viral spread rate on the binding rate? How does the reduction in binding rate impact plaque sizes in vitro experiments? Is there an optimal binding rate, which maximizes viral infection of the cell monolayer? Mathematical models in forms of reaction-diffusion equations for viral plaque size experiments have been used before in [30; 2; 39] and our model follows those principles.

To answer the above questions, we use three mathematical models that capture different aspects of viral dynamics. Model 1 focuses on a small-time scale (less than 16 hours), enabling observation of viral spread distances without considering cell death and viral replication. Model 2 extends the analysis to a longer time scale (about 5 days), incorporating viral spread, cell death, and viral replication. Model 3 then includes the cancer cells explicitly, which allows us to compare plaque sizes of different experiments. The results of our models exhibit excellent concordance with the observed experimental phenomena, providing valuable insights into the dynamics of reovirus-mediated oncolytic therapy. Based on the modelling we are then able to compute the optimal binding rate that leads to the largest plaque size.

### 1.1. Summary of experimental observations of [12]

Our mathematical models will be fitted and validated from the data of [12], where we have access to the original data set. In [12], characteristics of T3wt and SV5 are empirically determined on monolayers of TUBO cells (spontaneously derived HER2/neu positive murine breast cancer cells) and L929 cells (tumorigenic mouse fibroblasts). Monolayers of TUBO and L929 cells are exposed to T3wt or SV5 to measure cell attachment, virus replication, cell killing, and the size of plaques produced over several rounds of infection and re-infection. Reovirus typically bind cells within an hour, enter in 3-4 hours, replicate exponentially until 15-18 hours, and newly-made viruses become released from cells at 18-20 hours post-infection (hpi). Accordingly, short-term infections of 1 hour are used to monitor binding. Long-term infections of up to 5 days are used to monitor plaque size. In [11] they find that among a variety of reovirus mutations, the variant SV5 (supervirus 5), which has five mutations in the virus genome, leads to the largest plaque sizes. Relative to T3wt, SV5 displays similar kinetics of replication in an infected cell, cell death of the infected cell, burst size (the titer of virus released from infected cells), and diffusion. However, SV5 bindings less efficiently to cells, and also produces significantly larger plaques over several rounds of infection. While our mathematical modelling will use the empirical data derived on cancer cell cultures, it might be of interest to the reader that SV5 also significantly improves tumor regression and mouse survival in the more-complex mouse models of TUBO-derived tumors.

### 1.2. Outline

The paper introduces a multi-scale mathematical modeling approach to investigate the dynamics of viral spread in cell culture experiments. The study begins with a short time scale model in Section 2, utilizing data from [12] to estimate the binding rate *γ*_*b*_ and viral diffusion coefficient *D*_V_. Section 3 extends the model to longer time scales, incorporating events such as cell death and viral production. The speed of the spread of viral infection over the monolayer is computed via a travelling wave analysis, revealing the relationship between the binding rate *γ*_*b*_ and the viral spread speed *c*^∗^ in Section 3.6. Model validation is presented in Section 3.7, where additional experimental data is considered. In Section 3.8 and Section 4, we present the main result on the optimal choice of the binding rate *γ*_*b*_. The paper concludes in Section 5, contextualizing the results within the broader scope of mathematical modeling, and cancer and viral infection research.

## 2. Model 1: Short Time Scale

First, we start by answering the following question: How far does the virus spread depending on the binding rate during a time that is short enough to exclude effects of long-term cell death and viral replication? To answer this question, we use two sets of data from [12]. In short-time experiments, the percentage of virus binding to L929 cells after 1 hour was measured (see Table 1). These data will be used to estimate the binding rate *γ*_*b*_. Second, to estimate viral diffusion in the extracellular medium, the viral load was measured after inoculation in a cell free medium (see Figure 1).

**Table 1:**
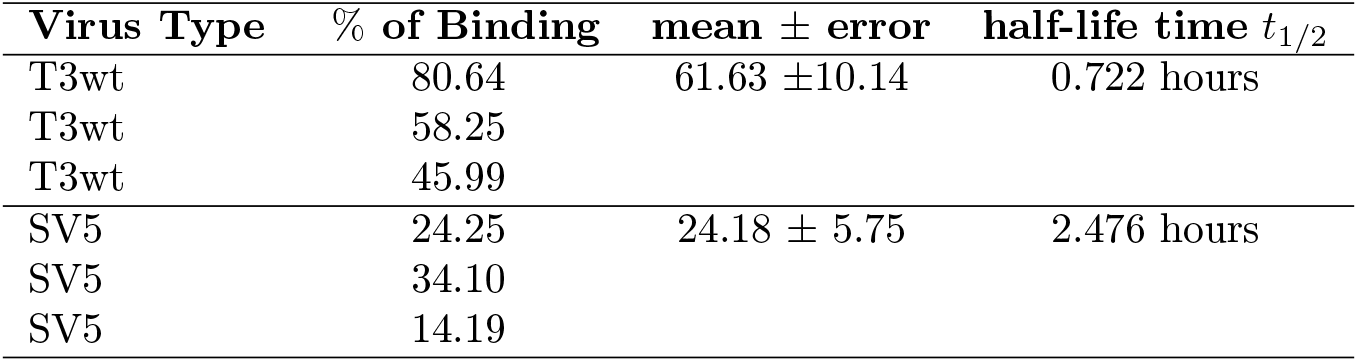
The percentage of binding for each virus type at 1 hour for three data sets each; taken from the original data that were used for Figure 6 A in [12]. The estimated data range is computed.

**Figure 1.**
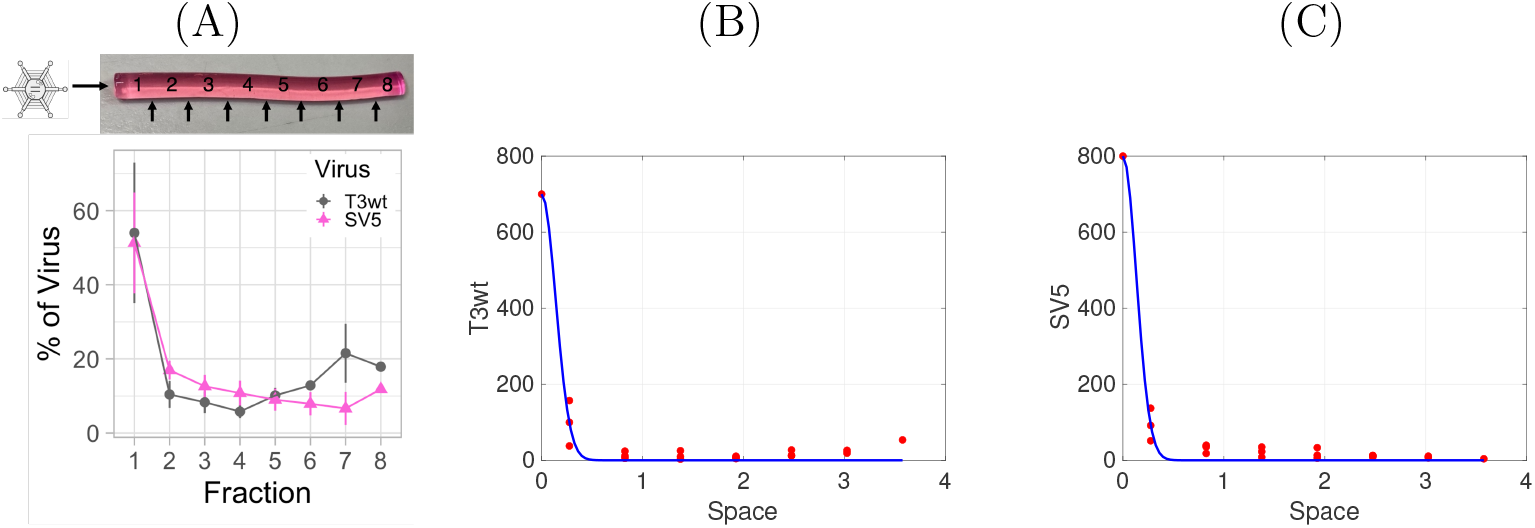
The diffusion coefficient estimation for T3wt and SV5 particles. **(A):** Percent of total T3wt and SV5 virus particles at each fraction measured following 120 hours of incubation [12], picture taken from Figure 7 A in [12]. **(B):** Plot of the original percentage data from [12] for T3wt and our fit as solid line. **(C):** Plot of the original percentage data from [12] for SV5 and our fit as solid line.

On the short time scale of 1-16h, we consider only two processes, which is binding of the virus to the cells and diffusion of virus particles in the cell medium. Hence our model has the form

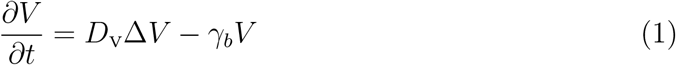

with the initial condition

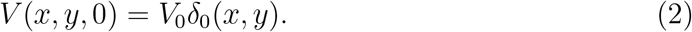

Here *V* (*x, t*) denotes the titre of virus particles, *D*_V_ is the constant diffusion coefficient, *γ*_*b*_ is the constant binding rate and *V*_0_ is the amount of virus particles at the start of the experiment *t* = 0. The symbol Δ denotes the Laplace operator, which is the sum of all second order partial derivatives, and *δ*_0_(*x, y*) denotes the delta distribution, which indicates the point of viral injection at the beginning of the experiment. We use model (1) to estimate the binding rate *γ*_*b*_ and the diffusion coefficient *D*_V_.

### 2.1. Binding Rate Estimation

To measure the binding rate *γ*_*b*_, we use the data in [12], where three experiments for each reovirus type were completed to estimate the efficacy of reovirus attachments to tumor cells. L929 cells were exposed to equivalent virus particle doses and incubated at 4^*o*^*C* for 1 hour to enable virus attachment without entry into the cells (i.e entry requires temperatures above 19^*o*^*C*). The unbound virus particles were removed by washing the cells extensively before harvesting the post-binding lysates. Finally, Western blot analysis was used to calculate the percentage of cell-bound virus particles based on virus protein levels in the lysates versus the input. The results show that on average 62% of T3wt virus were bound to L929 cells, compared to on average 24% of SV5 virus particles bound to cells (Table 1).

The binding process can be easily described by a linear binding law

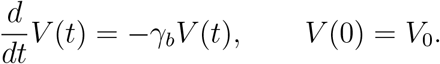

To estimate the binding rate *γ*_*b*_, we assume that the data in Table 1 are normally distributed and apply the likelihood method with the least square error (LSE) [16]. We denote the measured %-values of viruses binding as *y*_*i*_, for i=1,2,3, as there are three independent data points for each virus. The solution to the above equation is an exponential, which we can use to compute the number of unbound virus particles at time *t* as

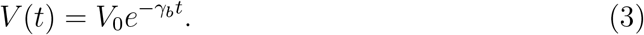

Then, the percentage of bound virus particles after *t* = 1 hour is

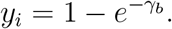

Therefore, *γ*_*b*_ = −ln(1 −*y*_*i*_). As there are several measurements for *y*_*i*_, an average is necessary, which is generated using the maximum likelihood estimator.

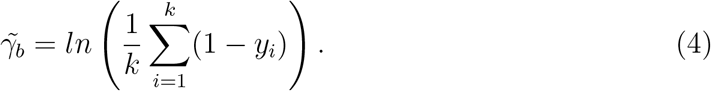

The corresponding confidence interval is then computed (see [16]). This estimation of the binding rate finds for the wild type (T3wt) 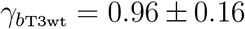 per hour, while for SV5 it is 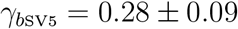 per hour.

These values can be related to the half-life of the virus population by

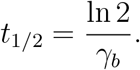

The T3wt virus population has a shorter time for binding with *t*_1*/*2_ = 0.722 hours compared to the SV5 virus population with *t*_1*/*2_ = 2.476 hours. Our results are consistent with the overserved measurements, as the data results indicated that T3wt virus particles have a higher percentage of binding with more than 50% of them binding compared to SV5 in one hour.

### 2.2. Diffusion Coefficient Estimation

The spatial diffusion of virus inside the culture medium without host cells was measured using the barrel of 1 mL syringes filled with semi-solid 0.5 % agar medium. The virus was introduced at the top of the medium and allowed to diffuse over time. The medium then removed from syringes and divided spatially into equal fractions (see Figure 1 A). Each fraction corresponds to 100 *μ* L. The percentage of viral load that diffused into each fraction was measured at time *t* = 120 hours. The spatial extent of each fraction is approximately 0.55 cm as we can see in Figure 1 A. To estimate the diffusion coefficient *D*_V_ at t=120 hour, we fit the data in Matlab by applying the Gaussian distribution formula for a diffusion process.

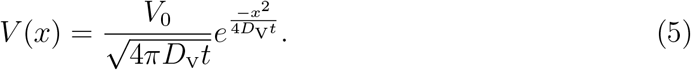

The parameters *V*_0_ and *D*_V_ are then estimated, where *V*_0_ is the number of viruses particles at t=0 which was not measured in the experiments. In Figure 1 we show the Figure 7 A from [12] in (A), and replot the original data with our fits in (B) and (C). We find that there were no significant differences in the diffusion coefficients between T3wt and SV5 viruses. The best fit estimated diffusion coefficient in both cases is *D*_V_ = 0.01 ±0.0015 mm^2^ per hour (equivalent *D*_V_ = 0.0001±0.000015 cm^2^ per hour) (Figure 1 B and C). The estimated values for *V*_0_ differ slightly, *V*_0,*T* 3*wt*_ = 243 and *V*_0,*SV* 5_ = 265, but they carry no biological information, as they are used to calibrate the distributions.

Another way to evaluate the diffusion coefficient *D*_V_ of small particles in a medium is by applying the Stokes-Einstein equation for the diffusion coefficient *D*_V_ of a spherical particle of radius *r* in a fluid of dynamic viscosity *η* at absolute temperature *T* [49]. We do not have any direct information from the data to estimate the value of the viscosity of the 0.5 % agar in Minimum Essential Media (MEM). Therefore, we use the viscosity of water which is also similar to the viscosity of Dulbecco’s Modified Eagle Medium (DMEM) (10 % FBS where FBS refers to a Fetal Bovine Serum) [23]. Thus, in our case the Stokes-Einstein relation gives

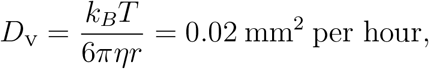

where *k*_*B*_ is Boltzmann’s constant, *r* = 35nm [33] is a typical virus radius, and *η* = 0.001 pa.s is the viscosity of water at *T* = 21^*o*^*C*. The values of diffusion coefficient in our estimation and Stokes-Einstein are very close. Furthermore, additional studies [15], [51] use a similar diffusion coefficient of *D*_Rioja_ = 0.014 mm^2^ per hour and *D*_P_ = 0.01 mm^2^ per hour for cancer viral therapy, which are similar to our value *D*_V_ = 0.01 mm^2^ per hour as we can see in Table 2.

**Table 2:** Comparison between the value of the viral diffusion coefficient estimated in our model versus the Stokes-Einstein relation, by the Rioja et.al model [15], and the Poolandvand et al. model [51].

| | $D_V$ (mm<sup>2</sup> per hour) |
| --- | --- |
| Our model fit estimation | $0.01 \pm 0.0015$ |
| Stokes-Einstein estimation [49] | 0.02 |
| Rioja et al. [15] estimation | 0.014 |
| Poolandvand et al. [51] estimation | 0.01 |

### 2.3. Result: Prediction of the Spread Radius for Short Times

Based on the above modelling and parameters we can estimate the distance a viral inoculation should spread on a short time scale. We assume the inoculation is somewhere in the centre of the monolayer such that viral spread is essentially radially symmetric. We consider the spread radius as the maximum distance from the inoculation after which no virus particles are detectable. The critical detection threshold for virus titer is denoted as *V*_min_. Hence we ask the question, for which radius *r* is the virus titer equal to *V*_min_?

For this we solve equations (1)-(2) with a little trick by setting 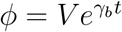. Then

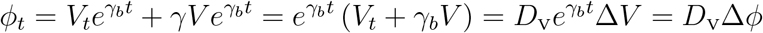

Hence *ϕ* satisfies a linear heat equation, which we can solve explicitly using the fundamental solution in 2-D [16].

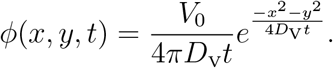

Using 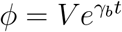, we obtain

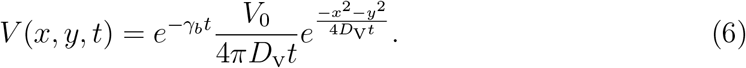

To compute the spread radius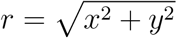, we assume that below a level of *V*_min_ no virus can be measured. Hence at the spread radius *r* we have

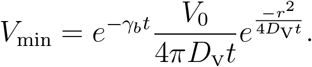

We solve this equation for *r* and obtain

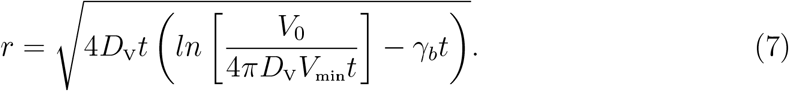

Fixing the time *t*, the diffusion coefficient *D*_V_, the number of virus particles at t=0 i.e *V*_0_, and the threshold *V*_min_, we have the spread radius *r* as a function of the binding rate *γ*_*b*_. We show this dependence in Figure 2. In Figure 2 we also show the estimated binding rate for the wild type virus in blue (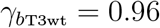 per hour) and for SV5 in red (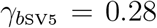 per hour). Furthermore, we see that the spread radius declines as the binding rate increases. For binding rates larger or equal 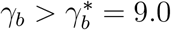, no more spread is possible, since all virus particles get bound to cells immediately.

**Figure 2.**
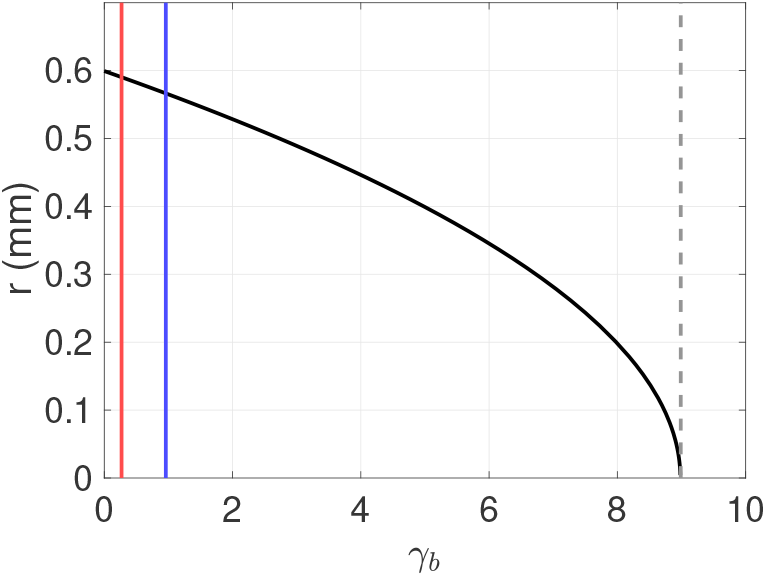
The plaque radius values as function of the binding rate *γ*_*b*_ (per hour) with *D*_V_ = 0.01 mm^2^ per hour, and 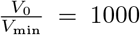 virus at t=1 hour. The blue line represents the value of *γ*_*b*T3wt_ = 0.96, while the red line represents the value of *γ*_*b*SV5_ = 0.28.

## 3. Model 2: Long Time Scale

### 3.1. Basic Assumptions and the Mathematical Model

In this section, we study the spread distances of the viral infection for time scales that include viral replication inside the cells, virion release and cell death (i.e. more than 16 hours). Specifically, we explain the plaque size results on L929 cell monolayers in [12]. Plaque size refers to the area of dead cells that result from the viral infection. The larger the plaque size, the farther the viral infection has spread. In [12], a monolayer of L929 cells was subjected to infection by reovirus particles. Following a one-hour incubation at 37^*o*^*C*, a 0.5% agar overlay was introduced onto the cells. Once the agar solidified, the cells were placed back into the 37^*o*^*C* incubator for a period of 5 days. Afterward, the cells were treated with 4% paraformaldehyde (PFA) for fixation and the cellular monolayer was stained using a 1% (wt/vol) crystal violet solution.

Subsequently, plaque size analysis was performed using the Fiji software with the particle analysis plugin and the results expressed as a relative plaque size to T3wt after normalization T3wt plaque size to 1. A larger plaque size reflects increased efficiencies of one or more steps involved in progressive infection and killing of more and more cells over cycles of virus infection, release, and reinfection.

To analyze the results of plaque size in [12], we assume that the number of cells that can be infected during the experiment is about constant. This is a strong assumption, but we feel that it is justified, since the model shows good results. We include a class of infected cells *I*(*x, t*) and extend our previous model (1) as

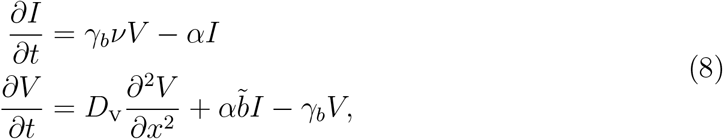

where the virus diffusion coefficient *D*_V_ and the binding rate *γ*_*b*_ are the same as before in model (1)-(2). The percentage of binding viruses that lead to infection is denoted by *ν*. The infected cells die at rate *α*, and the burst size of the infectious viruses is denoted by 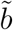. Here, we would like to indicate that the rate of virus replication in infected cell are expressed in some papers by parameter *b*, where *b* represents the infected cells death rate the × virus burst size [2; 13; 36]. This is equivalent in our model to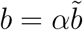.

In addition to *γ*_*b*_ and *D*_V_, we have three more parameters to estimate: the death rate of infected cells *α*, the percentage of binding virus that lead to an infection *ν*, and the burst size 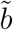. We estimate these values in the next subsections and summarize the values in Table 3.

**Table 3:** Estimated parameter values of system (8).

| Parameter | Description | Value | Unit |
| --- | --- | --- | --- |
| $D_V$ | Diffusion coefficient of reovirus | $0.01 \pm 0.0015$ | $\text{mm}^2$ per hour |
| $\gamma_{bT3wt}$ | Binding rate of wild type | $0.96 \pm 0.16$ | per hour |
| $\gamma_{bSV5}$ | Binding rate of SV5 | $0.28 \pm 0.09$ | per hour |
| $\alpha$ | Death rate of infected cells | $0.057 \pm 0.030$ | per hour |
| $\nu$ | Percentage of binding reovirus leads to infection | 0.01 | cell per virus |
| $\tilde{b}$ | Infectious burst size of reovirus | 500-1000 | virus per cell |
| $\tilde{b}_{T3wt}$ | Infectious burst size of T3wt | $514 \pm 114$ | viable virions per cell |
| $\tilde{b}_{SV5}$ | Infectious burst size of SV5 | $732 \pm 146$ | viable virions per cell |

### 3.2. Death Rate of Infected Cells Estimation

In [12] the percentage of cell death at time points 15, 18, 24, 30 and 36 hour after inoculation had been measured for each virus type T3wt and SV5. We show these data in Table 4 and illustrate them in Figure 3. While for 15 and 18 hours, only one data point had been measured compared to three data points at time 24 hour and two data points at time 30 and 36 hours each. From the data, we can see that the cells are surviving between 15-24 hours with no significant differences based on the virus type. The death rate of infected cells *α* at different time points can be estimated by data fitting of the exponential function *I*(*t*) = 1 −*e*^−*αt*^ using MATLAB, as shown in Figure 3 (B) and (C). We find no significant difference between the death rate of the T3wt virus and SV5 virus, which is consistent with the data observation. Therefore, we estimate the death rate of infected cells as *α* = 0.057 ± 0.030 per hour.

**Table 4:** The percentage of cell death at different time points for T3wt and SV5 particles from the experiment in Cristi et al. [12]. The estimated data range is computed.

| Virus Type | Time | % of Cell Death | Estimated Data |
| --- | --- | --- | --- |
| T3wt | 15 | 1.470 |  |
| T3wt | 18 | 40.15 |  |
| T3wt | 24 | 34.38 | $36.72 \pm 5.27$ |
| T3wt | 24 | 46.79 |  |
| T3wt | 24 | 28.99 |  |
| T3wt | 30 | 58.64 | $68.36 \pm 9.72$ |
| T3wt | 30 | 78.08 |  |
| T3wt | 36 | 62.35 | $71.49 \pm 9.14$ |
| T3wt | 36 | 80.63 |  |
| SV5 | 15 | 1.34 |  |
| SV5 | 18 | 39.05 |  |
| SV5 | 24 | 25.68 | $31.25 \pm 6.751$ |
| SV5 | 24 | 44.69 |  |
| SV5 | 24 | 23.39 |  |
| SV5 | 30 | 45.34 | $60.81 \pm 15.47$ |
| SV5 | 30 | 76.28 |  |
| SV5 | 36 | 52.15 | $64.99 \pm 12.84$ |
| SV5 | 36 | 77.83 |  |

**Figure 3.**
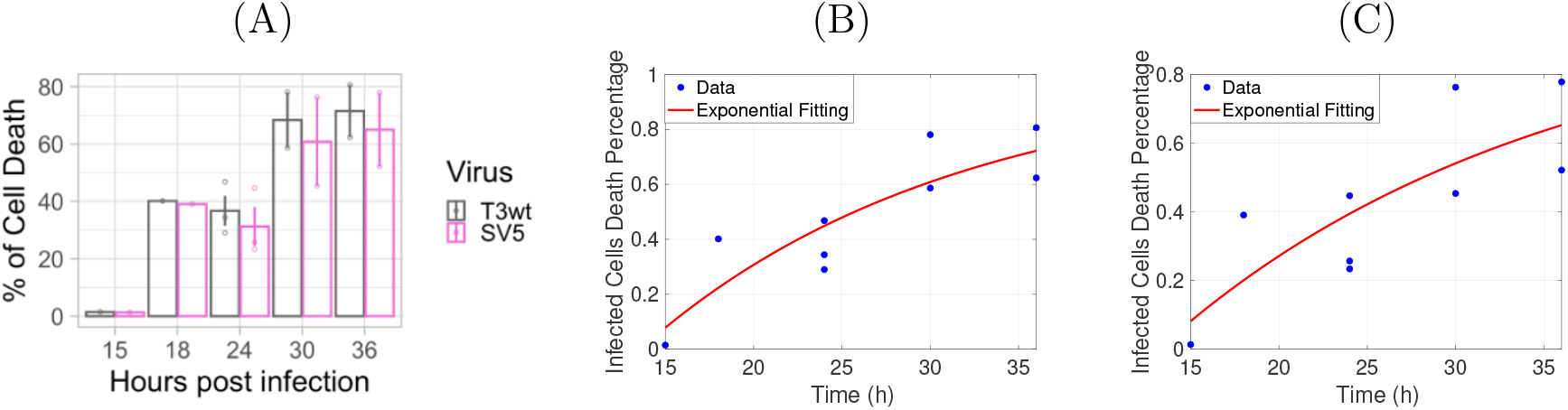
The death rate of infected cells estimation for T3wt and SV5 particles from experiment in Cristi et al. [12]. **(A)**: The percentage of cell death at different time points, picture taken from Figure 6 G in [12]. **(B)+(C)**: Replot of the original cell death data from [12] for T3wt in (B) and SV5 in (C) and our fits *I*(*t*) = 1 − *e*^−*αt*^ as red solid lines.

### 3.3. Viral Burst Size Estimation

The burst size of the virus is the number of released new virions from one infected cell. Experimentally in [12], the released number of virions had been measured with two data sets with multiplicity of infection (MOI) 21 for T3wt and 27 for SV5, respectively, at different time points: 0, 3, 6, 9, 12, 15, 18, 24, 30, and 36 hour. The Multiplicity of Infection (MOI) refers to the number of virions that are added per cell during infection. The data are shown in Figure 4. In Figures 4 (B), and (C) we use a logistic fit for these data and estimate the burst size as the carrying capacity value for this logistic fit. We find 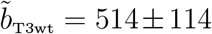 viable virions per cell and 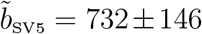 viable virions per cell, respectively. Note that the confidence intervals for these two values overlap. Hence, as reported already in [12], there is no statistically significant difference in those values. This is an important observation, and we come back to this issue later.

**Figure 4.**
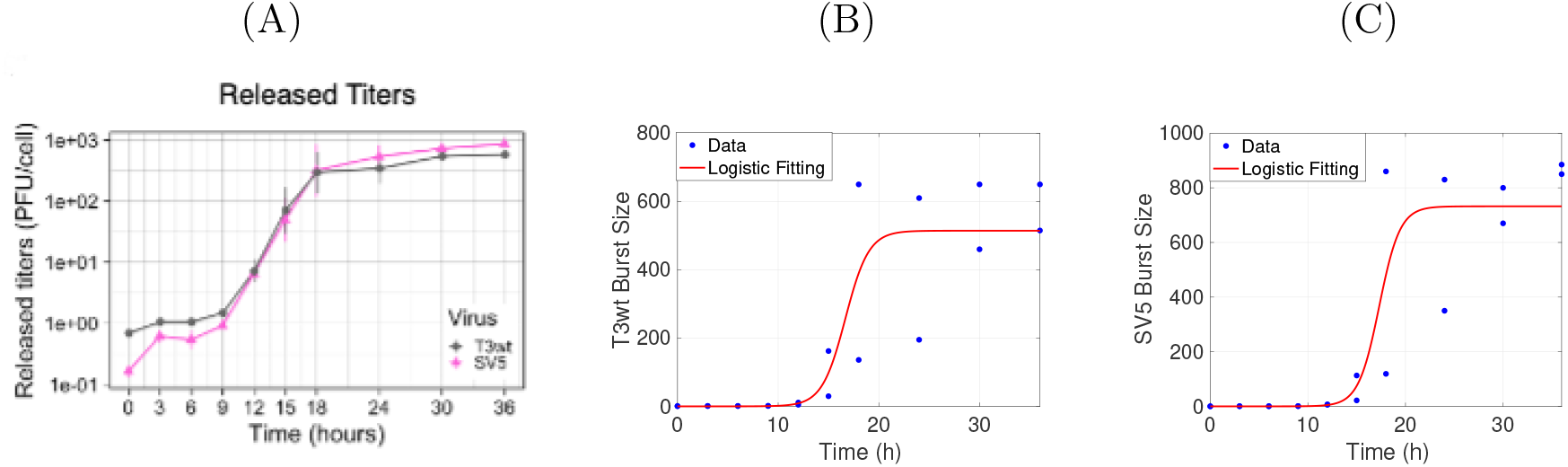
The burst size virus estimation for T3wt and SV5 particles from experiment in Cristi et al. [12]. **(A)**: The percentage of viral burst size at different time points, taken from Figure 6 F in [12]. **(B)+(C)**: Replot of the original burst size data from [12] for T3wt in (B) and SV5 in (C). Our logistic fits are shown as a red lines.

### 3.4. Percentage of Infectious Viral Particles

The results in [41], show that the percentage of bound reovirus that lead to an infection is in the range of 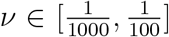 cells per virus. We choose *ν* = 0.01 cell per virus. This means out of 100 binding viruses on average one virus leads to a successful infection.

The parameters values are summarized in Table 3. Now, as all parameters for model (8) are identified, we can begin its analysis.

### 3.5. Viral Replication Number

A useful quantity for the analysis of our model is the virus replication number (VRN) i.e *R*_V_ of the ODE system (8). Defined in [2] as analogy of the basic reproduction number in epidemiology, the viral replication number denotes the average number of infected cells that result from one infected cell in an otherwise healthy cell population. The viral replication number is used as a measure to quantify the transmissibility in the cell culture. Since the system (8) is linear and the only steady state is (*I, V*) = (0, 0), then the corresponding eigenvalue problem is

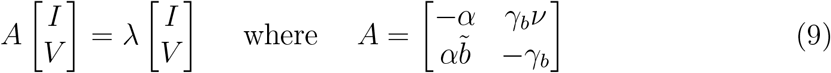

Condition det *A* = 0 can be written as *R*_V_ = 1, where

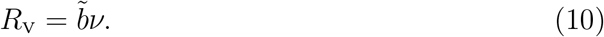

The stability of the steady state (*I, V*) = (0, 0) is determined by *R*_V_. Here, *R*_V_ denotes the average burst size. Mathematically, if *R*_V_ *<* 1, then both eigenvalues of the matrix A are negative and hence the steady state (0,0) is stable. Biologically, this means that the virus dies out. On the other hand, if *R*_V_ *>* 1, then we have one positive eigenvalue and the second eigenvalue is negative. Therefore, the steady state is unstable and hence the virus can spread to infect the neighbouring cells and as a result the virus population grows. From the experimental data, we find that 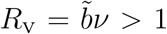, where *R*_VT3wt_ = 5.14 ± 1.14 and *R*_VSV5_ = 7.32 ± 1.46, respectively.

### 3.6. Travelling Invasion Wave

One way to understand the effect of the binding rate *γ*_*b*_ on the invasion of the viral infection over the L929 cell culture is an invasion wave analysis. This is a standard method for reaction-diffusion models [9] (like (8)) where an invasion speed is estimated, which describes the speed of the spread of the viral infection over the population of cells. Here we assume that *R*_V_ *>* 1, such that the virus can grow. Since our model (8) is linear we use the leading edge method, where we focus on the behaviour of the front profile of the invasion, near the steady state (*I, V*) = (0, 0). We use *c* to denote the invasion wave speed and *λ* to denote its exponential decay rate and look for solutions of the form

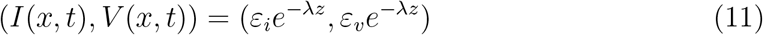

with *z* = *x* −*ct* and small contants *ε*_*i*_, *ε*_*v*_.

Substituting the Ansatz for (*I*(*x, t*), *V* (*x, t*)) into system (8), we get

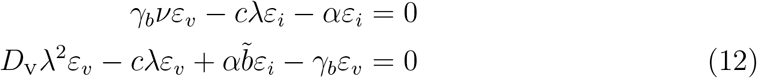

We write system (12) in matrix form *Aε* = 0,

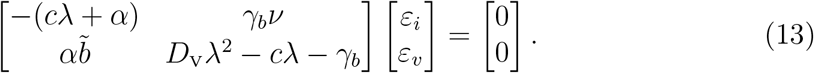

Thus, to obtain a non-trivial solution, we assume that the determinant of the matrix *A* is zero. The characteristic equation of (13) becomes

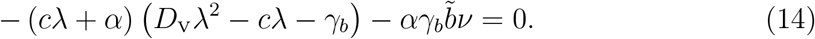

Following a method of Volpert [2], we introduce *ϱ* = *cλ >* 0, substitute it into (14) and solve for *c*^2^ to obtain

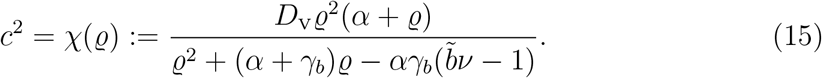

Next we show that *χ*(*ϱ*) has a unique positive minimum at *ϱ*^∗^ such that the minimal wave speed *c*^∗^ is given by

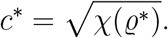

Consequently, we find the decay rate of the invasion with minimal speed as

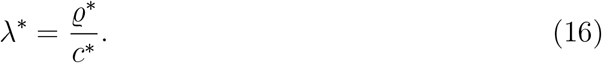

For the parameter values that we estimated for the two virus types T3wt and SV5, as reported in Table 3, we plot the curves *χ*(*ϱ*) in Figure 5 (A). The function *χ*(*ϱ*) has zeros at *ϱ* = 0, and *ϱ* = −*α*, which are not relevant since we require *ϱ >* 0. Also, *χ*(*ϱ*) has a vertical asymptote at

**Figure 5.**
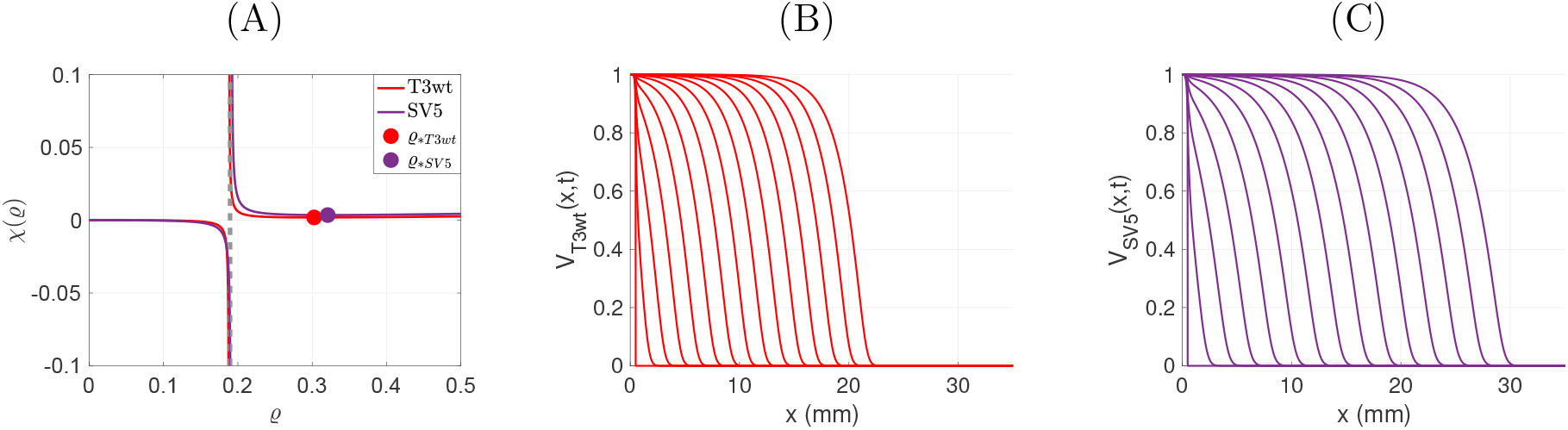
(A) Characteristic function *χ*(*ρ*) for the two cases of T3wt (red) and SV5 (purple) and the corresponding minima marked as solid points. (B) and (C) show the travelling wave as function of space, where the wave profiles for different time steps are overlayed. **(B)**: *γ*_*b*T3wt_ = 0.96, with 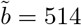, and *c* = 0.04373 mm per hour. **(C)**: *γ*_*b*SV5_ = 0.28, with 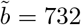 and *c* = 0.0599 mm per hour.

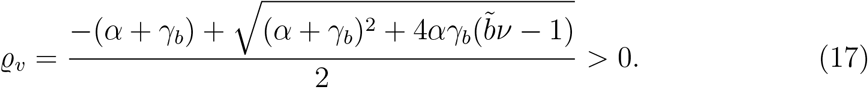

The positive minima, indicated as dots in Figure 5 (A), are right of the asymptote, hence we define

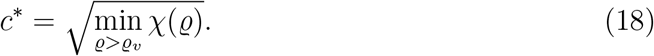

For the parameter values from Table 3 we get

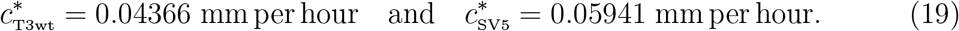

In Figure 5 (B) and (C) we show numerical simulations of the invasion waves for these parameters. We see that the invasion wave of T3wt (B) is slower than the invasion of the SV5 virus (C). If we compute the invasion speeds numerically, we find

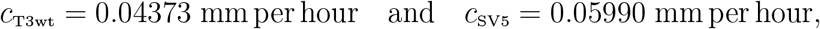

which is very close to the theoretical values above (19).

We also determine the corresponding invasion front decay rate (*λ*^∗^) by formula (16) and find

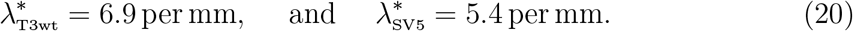

### 3.7. Validation of Model 2 on Invasion Front Data

In the previous sections we estimated all the model parameters as summarized in Table 3, plus the invasion speeds *c*^∗^ and the decay rates *λ*^∗^ at the edge of the invasion front. To validate our model (8) we compare it now to data that have not been used to parameterize the model. The set of data is an experiment in [12] where we use fluorescence measurements of viral load at the edge of the plaques.

When plaques are evaluated by crystal violet staining as in above experiments, then plaque size only reflects the size of clearance produced by killing of cells in the center. Crystal violet staining does not, however, reveal the extent of cells that are infected by virus but are still alive. Therefore, immunofluorescence was used to directly visualize reovirus-infected cells in the margin of the plaques at days 2, 3 and 4 post infection [12] (see Figure 6). In these experiments, SV5 infected cells were found at further distances from the origin than for T3wt. Mathematically, this can be represented by estimating the decay rate *λ* of the invasion front (11) of T3wt and SV5 viruses, which we considered earlier.

**Figure 6.**
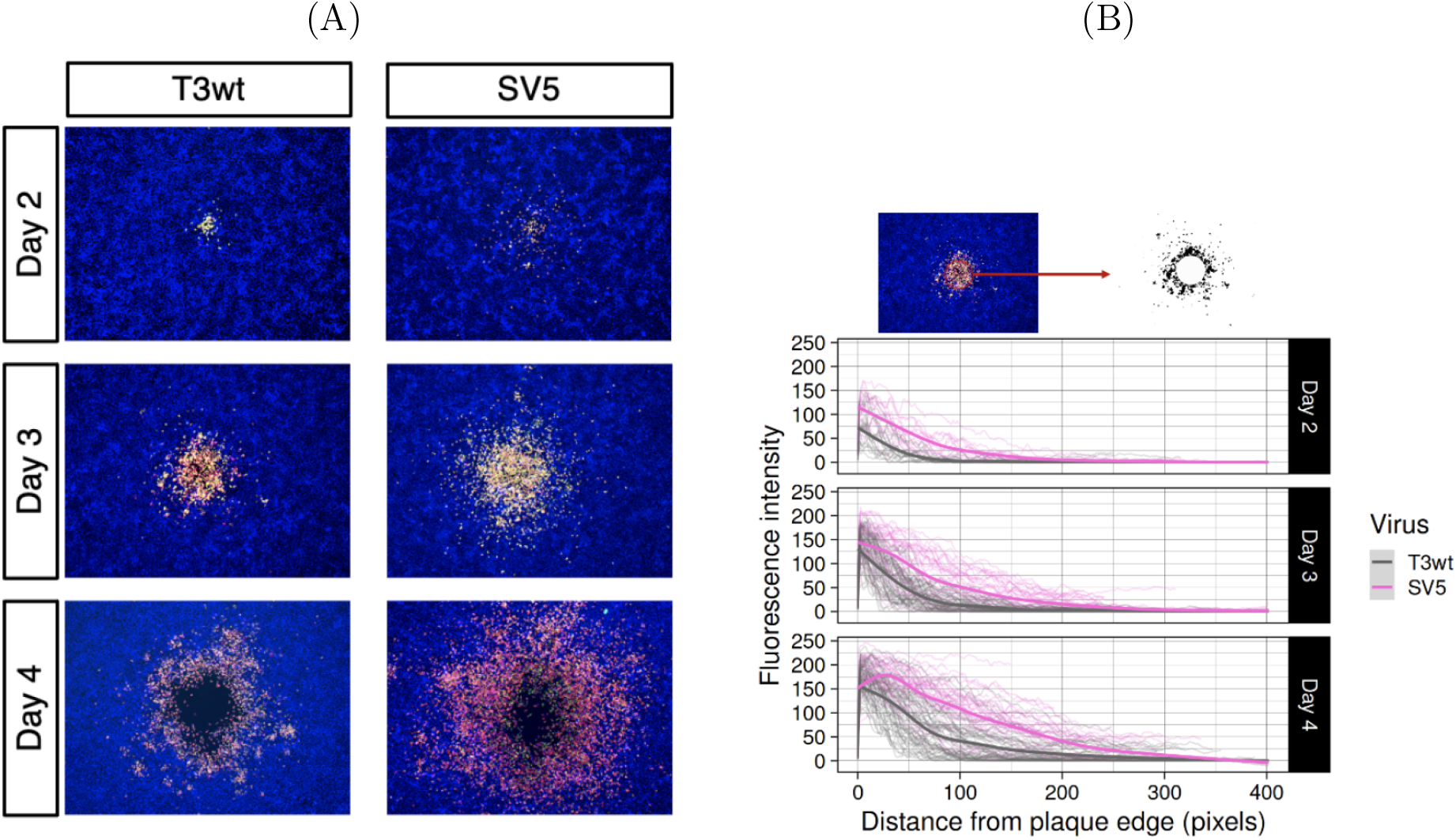
SV5 spread farther than T3wt. **(A)**: Immunofluorescence pictures of T3wt and SV5 plaques formation at days 2, 3 and 4 post infection from Figure 6 H in [12], **(B)**: Quantification of the fluorescence from the edge of the plaque to represent the spread of the virus from Figure 6 I in [12].

To estimate the decay rates of the T3wt invasion front *λ*_T3wt_ and the SV5 invasion front *λ*_SV5_ from the data, we apply MATLAB to fit the data in Figure 6 with an exponential decay function for the value *Fl*(*x, t*) = *e*^−*λ*(*x*−*s*)^, where *λ* is the invasion front decay rate and *s* is a shift of the exponential decay function to place it at the best location for the fit. In Figures 7 And 8 we show this fit in red with the corresponding data in blue. In Figure 7 we find the best fit decay rates on days 2, 3, 4 for T3wt to be *λ* = 0.029±0.001, 0.022 ± 0.0004, 0.014 ± 0.0003 per pixel, which has a mean value of *λ* = 0.022 per pixel. There are 445 pixel per mm, hence we find *λ*_T3wt_ = 9.8 per mm. This corresponds well with the previous estimate in (20) of 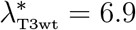 per mm. For the supervirus SV5 we find the decay rates *λ* = 0.015±0.0005, 0.011± 0.0002, 0.007 ±0.0002 per pixel, with an average of 0.011. This corresponds to *λ*_SV5_ = 4.9 per mm. Again, this is very close to the theoretical value of 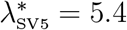 per mm from (20).

**Figure 7.**
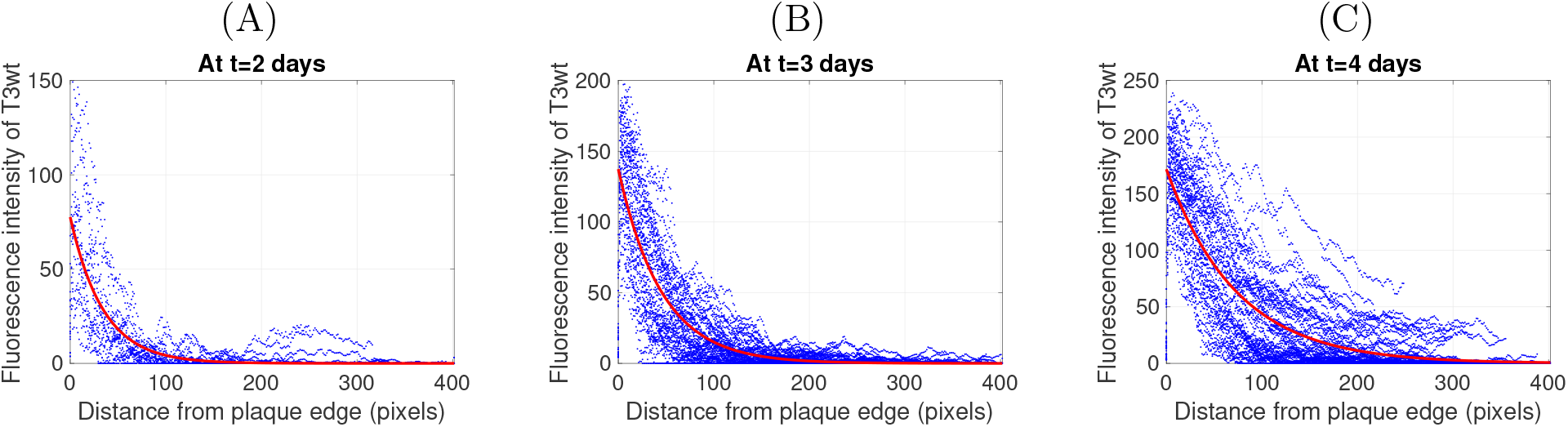
The T3wt invasion front decay rate. We plot in blue the original data from [12] and overlay the exponential decay function in red. **(A)**: *λ* = 0.029 at t= 2 days **(B)**:*λ* = 0.022 at t=3 days and **(C)**:*λ* = 0.014 at t=4 days.

**Figure 8.**
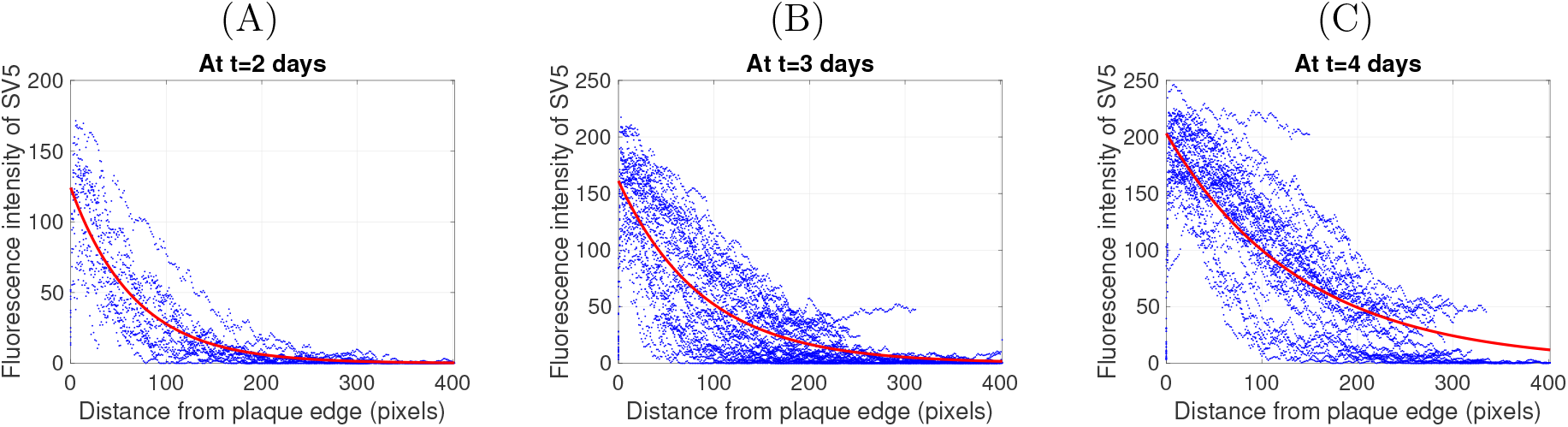
The SV5 invasion front decay rate. We plot in blue the original data from [12] and overlay the exponential decay function in red. **(A)**: *λ* = 0.015 at t= 2 days **(B)**: *λ* = 0.011 at t=3 days and **(C)**: *λ* = 0.007 at t=4 days.

### 3.8. The Relationship Between Binding Rate and Wave Speed

The relationship between the binding rate *γ*_*b*_ and the speed of the viral infection wave over the cell monolayer is very important. To optimize the efficacy of reovirus treatment, we like to find the binding rate 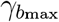 that maximizes the invasion speed *c*^∗^. A formula for *c*^∗^ is given in (18), where the function *χ*(*ρ*) and the value of the asymptote *ρ*_*v*_ both depend on the binding rate *γ*_*b*_. We first look at the extreme cases of no binding, *γ*_*b*_ = 0, and of immediate binding, *γ*_*b*_→ ∞.

In the case of no virus binding (*γ*_*b*_ = 0), there will be no invasion, since the virus cannot replicate. In this case we expect *c*^∗^ = 0. Indeed, in case of *γ*_*b*_ = 0, direct calculation in (15) and (17) shows that

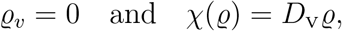

and

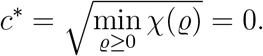

This means when the binding rate *γ*_*b*_ = 0, the invasion speed for T3wt and SV5 viruses infection is zero.

In the other extreme of immediate binding *γ*_*b*_→ ∞, the virus will bind immediately to cells and will no longer be able to invade any further. So we also expect *c*^∗^ = 0. In this case we have the singularity of *χ*(*ρ*) at 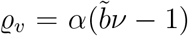. For *ρ > ρ*_*v*_ we can apply L’Hopital’s rule to *χ* given in (15) to consider the limit as *γ*_*b*_ → ∞. We find

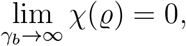

which implies *c*^∗^ = 0.

Since in the two extreme cases of *γ*_*b*_ the invasion speed is zero, and since *χ* and *ρ*_*v*_ depend continuously on *γ*_*b*_ for *ρ > ρ*_*v*_, we conclude that there is at least one maximum of *c*^∗^ for some intermediate binding rate *γ*_*b*_ ∈ (0, ∞). To find this value we consider the critical points of *χ*(*ρ*):

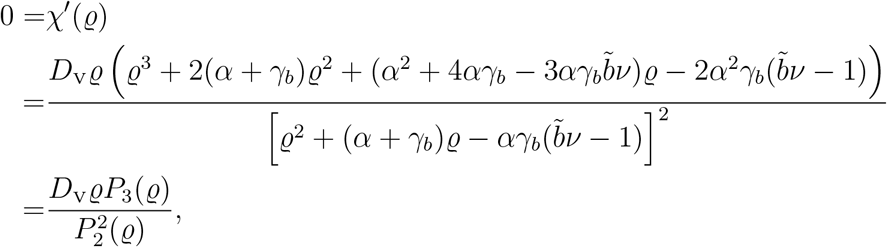

where

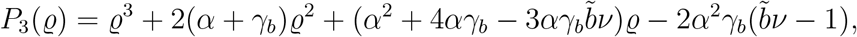

and

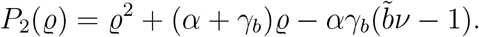

Thus, *χ*^′^(*ϱ*) = 0 when *ϱ* = 0 or *P*_3_(*ϱ*) = 0. Clearly, the coefficient of *ϱ*^3^ and *ϱ*^2^ are positive while the sign of the coefficient of *ϱ*^0^ is negative. The sign of the coefficient of *ϱ* depends on the value of *γ*_*b*_ after fixing the parameter values of 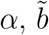, and *ν*. Based on Descartes’ rule of signs [8] even if the sign of the coefficient *ϱ* is positive or negative, we have only one positive real root and 2 or zero negative real roots of *P*_3_. Therefore, there exist a unique *ϱ*^∗^ *>* 0, such that *χ*^′^(*ϱ*^∗^) = 0. Furthermore, at *ϱ* = 0, we have 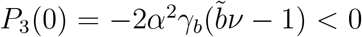, with the continuity of *P*_3_(*ϱ*) and being concave up since 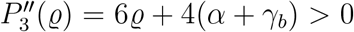 for each *ϱ*≥ 0. Therefore, by intermediate value theorem and mean value theorem, there is only one positive real root i.e *ϱ*^∗^ *>* 0 such that *P*_3_(*ϱ*^∗^) = 0. Thus for each *γ*_*b*_ *>* 0, *χ*(*ϱ*^∗^) is a unique minimum with *P*_3_(*ϱ*^∗^) = 0 for *ϱ*^∗^ *> ρ*_*v*_. The maximum possible invasion speed is then

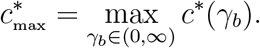

For the parameter values from Table 3 for T3wt and SV5 we plot the function *c*^∗^(*γ*_*b*_) as red line for T3wt and in purple for SV5 in Figure 9 (A). As red and purple points we indicate the estimated binding rates from the data for the corresponding cases, and in black we indicate the maximum of these curves.

**Figure 9.**
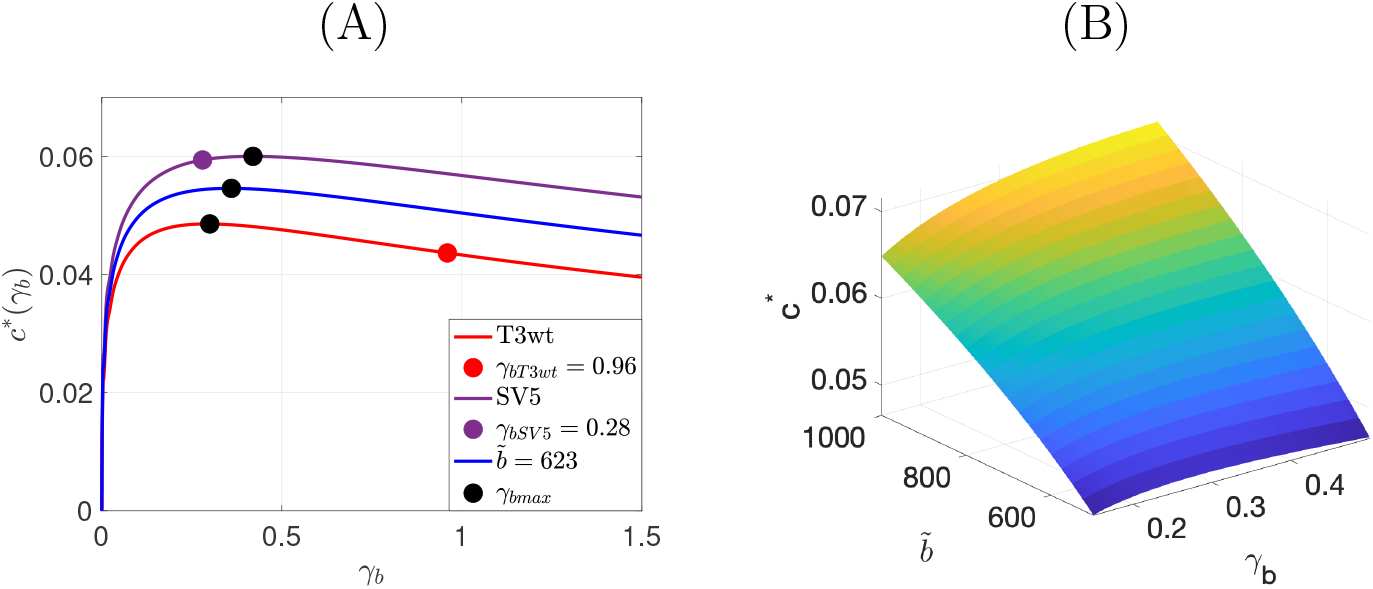
(A) The relationship between the binding rate *γ*_*b*_ and the invasion speed *c*^∗^ for T3wt, SV5, and when 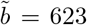. We plot the function (15) for different values of *γ*_*b*_ to find the maximum *γ*_*b*max_ that leads to maximum wave speed i.e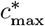. The numerical results indicate that for T3wt virus, the 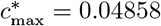 at *γ*_*b*max_ = 0.29, while the 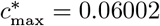 at *γ*_*b*max_ = 0.42 for SV5 virus. Finally, when we choose intermediate value of infectious burst size 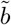, we find the 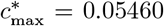 at *γ*_*b*max_ = 0.36. (B) The values of the minimum wave speed *c*^∗^ when we vary the binding rate *γ*_*b*_ and the burst size 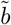. The maximum wave speed *c*^∗^ is 0.07154 when *γ*_*b*_ = 0.5 and 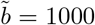. The values of the binding rate in (B) have the range *γ*_*b*_ ∈ [0.15, 0.5].

- For T3wt (red curve) we observe that the invasion speed could be increased by reducing the binding rate *γ*_*b*_ from 0.96 per hour to 0.29 per hour. In that case the speed would change from 0.044 mm per hour to 0.048 mm per hour. Expressed in percentage of binding after one hour, we aim to decrease the percentage of binding of T3wt virus from 61.7 % to 25.9 %.
- In the case of SV5 (purple curve) we see that an increase in binding rate from 0.28 to 0.48 would have a small accelerating effect from *c*^∗^ = 0.059 to 0.060. In other words, we like to increase the percentage of binding of SV5 virus after one hour from 24.4 % to 34.3 %.
- We also notice that this result depends on the burst size of the corresponding virus. As indicated earlier, and also in [12], the difference in the burst sizes is not statistically significant. Hence we add, in blue, the corresponding curve for the mean burst size of 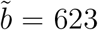. The curve is very similar to the red and purple curves and the maximum invasion speed of *c*^∗^ = 0.054 is found for a binding rate near 0.36.
- We also considered some extreme cases for the burst size of 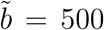 and 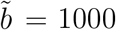. For 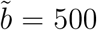 we find 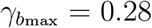 per hour with corresponding wave speed 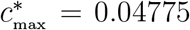 mm per hour, while for the upper bound of burst size 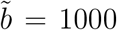 virus per cell we find 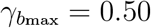 per hour with corresponding wave speed 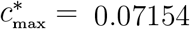 mm per hour.

The previous results emphasize the importance of the burst size parameter 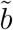 in determining the viral spread *c*^∗^ and, as a result, the plaque size. In Figure 9 (B), we determine the minimum wave speed values, denoted as *c*^∗^, by varying the binding rate *γ*_*b*_ and the burst size 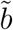. It is observed that for our range of possible burst sizes 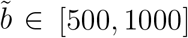, the maximum wave speed *c*^∗^ reaches 0.07154 when *γ*_*b*_ = 0.5 and 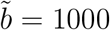.

We observe that the ratio 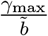 in Table 5 and Table 6 remains nearly constant for each maximum binding rate *γ*_max_ corresponding to the burst size 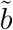. Hence, we might use the ratio 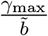 as a benchmark to assess how closely the experimental results approach the maximum required viral spread speed. The average number of ratio 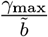 is 0.00056. Therefore, the optimal binding rate and burst size have a ratio of about 0.00056.

**Table 5:** The infectious burst size, the binding rate, % of binding viruses and the corresponding wave speed for the wild type T3wt, intermediate infectious burst size, and SV5 virus.

| | $\gamma_b$ | % of Binding | $c^*$ | $\tilde{b}$ | $\frac{\gamma_{\max}}{\tilde{b}}$ |
| --- | --- | --- | --- | --- | --- |
| T3wt Data | 0.96 | 61.7 | 0.04366 | 514 |  |
| Max T3wt | 0.29 | 25.2 | 0.04858 | 514 | 0.00056 |
| Intermediate Max | 0.36 | 30.2 | 0.05460 | 623 | 0.00058 |
| SV5 Data | 0.28 | 24.4 | 0.05941 | 732 |  |
| Max SV5 | 0.42 | 34.3 | 0.06002 | 732 | 0.00057 |

**Table 6:** The ratio of infectious burst size, the corresponding maximum binding rate, the maximum wave speed and the corresponding plaque size. The plaque size is computed with Model 3 as described in Section 4.

| $\tilde{b}$ | $\gamma_{\text{bmax}}$ | $\frac{\gamma_{\text{bmax}}}{\tilde{b}}$ | $c_{\text{max}}^*$ | Plaque Size |
| --- | --- | --- | --- | --- |
| virus per cell | per hour | cell per (virus × hour) | mm per hour | mm<sup>2</sup> |
| 500 | 0.28 | 0.00056 | 0.04775 | 5 |
| 550 | 0.32 | 0.00058 | 0.05065 | 20 |
| 600 | 0.36 | 0.00060 | 0.05338 | 32 |
| 650 | 0.36 | 0.00055 | 0.05600 | 43 |
| 700 | 0.39 | 0.00056 | 0.05848 | 55 |
| 750 | 0.43 | 0.00057 | 0.06090 | 69 |
| 800 | 0.47 | 0.00059 | 0.06316 | 82 |
| 850 | 0.47 | 0.00055 | 0.06538 | 95 |
| 900 | 0.51 | 0.00057 | 0.06753 | 106 |
| 950 | 0.51 | 0.00054 | 0.06959 | 121 |
| 1000 | 0.50 | 0.00050 | 0.07154 | 129 |

## 4. Model 3: Plaque Size

### 4.1. Plaque Size Experiments

In [12] an experiment is designed to measure the plaque size of the T3wt and SV5 viruses. They reported the relative areas *A*_SV5_*/A*_T3wt_ and found that the relative value of plaque size between SV5 and T3wt after 5 days varies between 3.4530 to 5.1248. This means that after 5 days, the plaque size of SV5 virus is about 4 times larger than the plaque size of the T3wt virus. We would like to point out the reason to measure the relative value of plaque size. It was observed that repeat experiments lead to different plaque sizes, due to variables that are out of control of the experimentalist such as cell viability, humidity, person performing the experiments, etc.. However, the relative plaque size difference of a factor of 4 were similar in all experiments. The reported values for *A*_SV5_*/A*_T3wt_ are

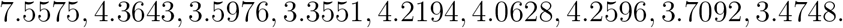

with mean and standard error

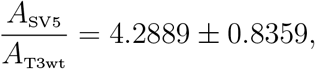

which we like to confirm with our model.

To properly keep track of the plaque sizes, we now include the cancer cell compartment *C*(*x, t*) explicitly. The plaques correspond to regions of dead cancer cells, and in our modelling we identify those as regions where *C*(*x, t*) is below a small threshold. Our previous model (8) is now extended to **Model 3:**

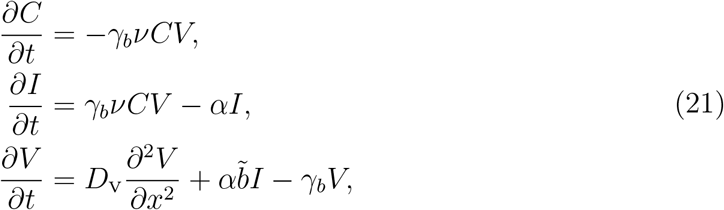

where all parameters have already been identified in Table 3.

We numerically solve our model (21) with virus inoculated in the center of a two-dimensional domain (see Figure 10). We estimate the plaque sizes after 5 days with threshold for cancer cells of 1%, indicated as a red line in the figures. We find a ratio 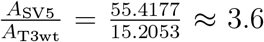, which is very close to the experimental ratio mentioned above.

**Figure 10.**
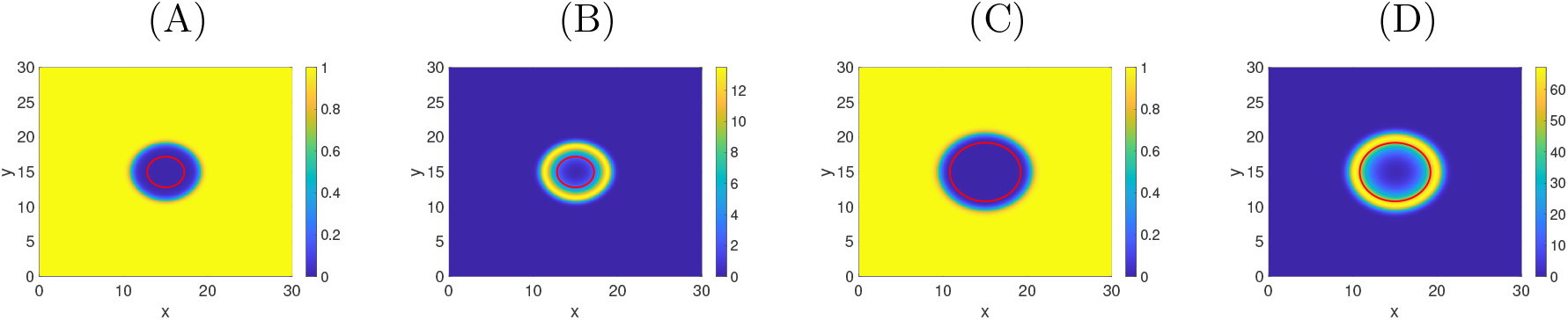
The plaque sizes at 5 days. **(A)+(B)**: T3wt spread with cancer cell density in (A) and viral concentration in (B). The *C*-level of 1% is indicated as a red line. **(C)+(D)**: SV5 infection with cancer cells in (C) and SV5 in (D). The computed plaque sizes are indicated in the red circle in (A) which is 15.2053 for T3wt virus and 55.4177 for SV5 virus in (C). Therefore, the relative plaque size of SV5 related to T3wt is 3.6446.

We note that the T3wt and SV5 invasion forms a hollow ring spread pattern (see Figure 10). Such invasion patterns are typical for virus infections of tissues, and were also previously found in [61; 4].

Furthermore, we perform simulations of this model for a few chosen parameter values to see the dependence on *γ*_*b*_ and 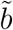. In Figure 11 And 12, we fix all parameters as in Table 3, while the burst size 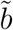 is varied as follows: 514 (T3wt), 623 (average), 732 (SV5), and 1000 (max). A notable increase in plaque size is observed for increased burst size.

**Figure 11.**
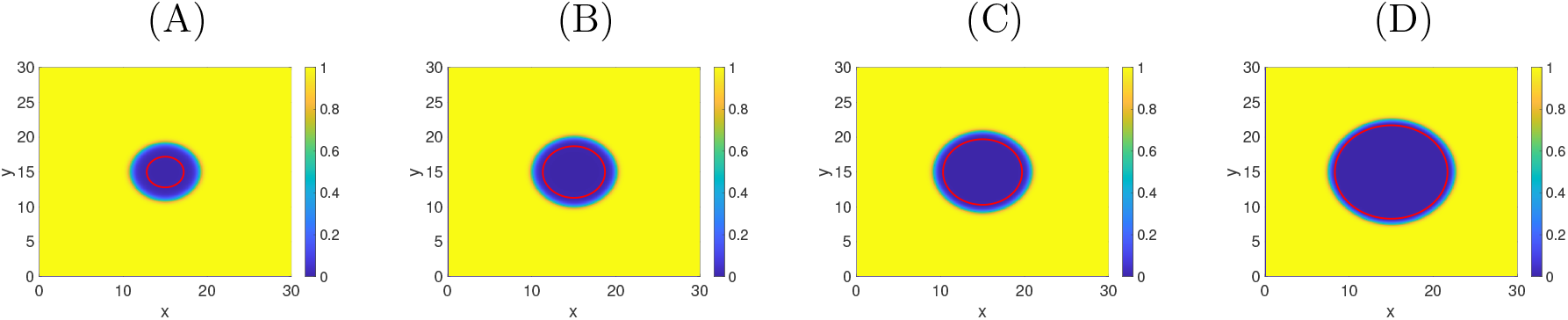
The plaque sizes of T3wt at t=5 days when *γ*_*b*_ = 0.96 with different burst sizes 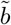. **(A)**: 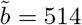 **(B)**: 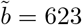**(C)**: 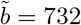 and **(D)**: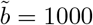. The threshold= 1 %.

**Figure 12.**
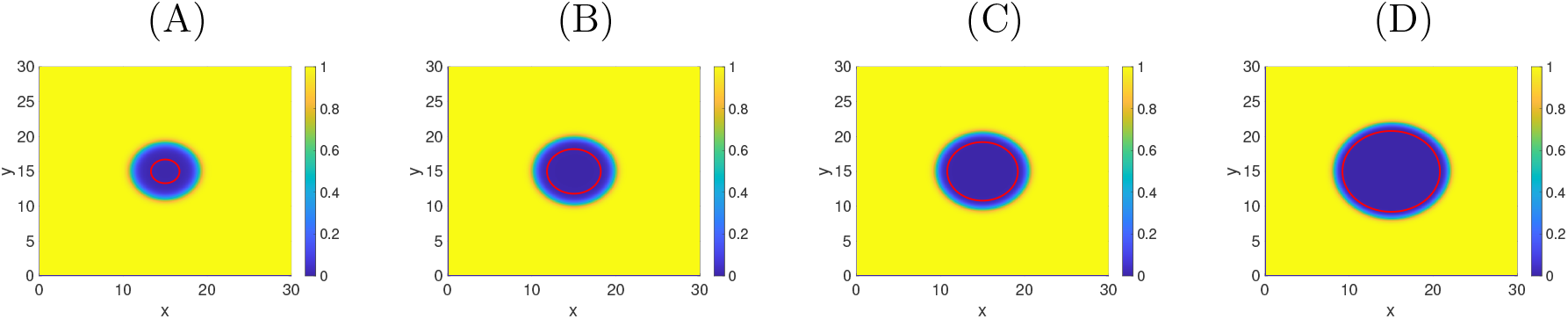
The plaque sizes of SV5 at t=5 days when *γ*_*b*_ = 0.28 with different burst size 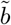. **(A)**: 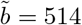 **(B)**: 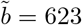 **(C)**: 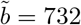 and **(D)**: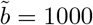. The threshold=1 %.

The paper [12] presents an additional dataset that we have not included here. These data pertain to plaque experiments conducted under the administration of the drug neuraminidase. Neuraminidase, known as a cancer chemotherapy agent, reduces the binding affinity of the virus. Our extended model (21) appears to offer the appropriate level of detail to simulate these experiments, and we are currently engaged in discussions regarding the specifics of this modeling.

## 5. Conclusion

This study employs a reaction-diffusion model to investigate key aspects of viral infection dynamics of cancer cell monolayers. Specifically, we explore the impact of the binding rate on the spread of the viral infection over the monolayer, the correlation between viral invasion speed and binding rate, and the repercussions of reducing binding rate on plaque size. Two distinct time scales are considered: a short duration (less than 16 hours) focusing on viral spread preceding cell death and replication events, and a longer time scale addressing viral infection between cells. All the parameters in our models are estimated using data from [12].

Our model establishes that the maximum speed of viral infection aligns with a fine balance of viral binding and burst size. The binding has to be fast enough to allow for efficient cell infection, but in the absence of fast flowing medium, it also has to be weak enough to allow the virus to spread longer distances before binding. These observations from the mathematical modelling contradict the common belief that increased binding rate leads to increased viral infection. If binding is increased too much, the viral particles get trapped locally and cannot invade into the remaining tissue. For the specific case of reovirus considered here, our findings indicate that 25.9% binding after one hour is optimal for T3wt virus to achieve the maximum viral spread rate, whereas the SV5 virus infection is optimized for approximately 34.3% binding after one hour. Based on these findings, reovirus mutant can now be screened for binding efficiency with a specific objective to find a mutant that can best fulfil this postulated optimum. In previous studies, reovirus plaque size on cancer cells in vitro correlated with significantly improved oncolytic activity in vivo. We can predict therefore that a reovirus mutant of burst size 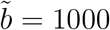 with 39.3 % binding would not only generate larger plaques than T3wt on cancer cells in vitro, but reduce tumor size and improve survival also relative to T3wt in mouse models of breast cancer.

## CRediT authorship contribution statement

**Arwa Abdulla Baabdulla:** Formal analysis; investigation; methodology; software; validation; visualization; writing – original draft; writing – review and editing.

**Francisca Cristi:** Data curation.

**Maya Shmulevitz:** Conceptualization; data curation; review and editing.

**Thomas Hillen:** Conceptualization; methodology; supervision; writing – original draft; writing – review and editing.

## Acknowledgements

AAB acknowledges support through United Arab Emirates University Scholarship. TH acknowledges support from the Natural Science and Engineering Research Council of Canada (NSERC). AAB and TH thank the members of the Mathematical Biology Journal Club for their valuable comments. MS acknowledges support from the Cancer Research Society (CRS) and the Canadian Institutes of Health Research (CIHR), as well as salary support from the Canada Research Chairs (CRC) and infrastructure support from the Canada Foundation for Innovation (CFI). FC acknowledges scholarship support from the John and Rose McAllister Graduate Scholarship award from the Faculty of Graduate Studies and Research at the University of Alberta, the Faculty of Medicine & Dentistry (FoMD) Dean’s Doctoral Award from the University of Alberta, the LKSIoV Doctoral Award, and the La Vie en Rose Scholarship for Breast Cancer Research from the Cancer Research Institute of Northern Alberta (CRINA).

## Data availability

Data is available in publication [12].

